# Dissecting quantitative trait nucleotides by saturation genome editing

**DOI:** 10.1101/2024.02.02.577784

**Authors:** Kevin R. Roy, Justin D. Smith, Shengdi Li, Sibylle C. Vonesch, Michelle Nguyen, Wallace T. Burnett, Kevin M. Orsley, Cheng-Sheng Lee, James E. Haber, Robert P. St.Onge, Lars M. Steinmetz

## Abstract

Genome editing technologies have the potential to transform our understanding of how genetic variation gives rise to complex traits through the systematic engineering and phenotypic characterization of genetic variants. However, there has yet to be a system with sufficient efficiency, fidelity, and throughput to comprehensively identify causal variants at the genome scale. Here we explored the ability of templated CRISPR editing systems to install natural variants genome-wide in budding yeast. We optimized several approaches to enhance homology-directed repair (HDR) with donor DNA templates, including donor recruitment to target sites, single-stranded donor production by bacterial retrons, and in vivo plasmid assembly. We uncovered unique advantages of each system that we integrated into a single superior system named MAGESTIC 3.0. We used MAGESTIC 3.0 to dissect causal variants residing in 112 quantitative trait loci across 32 environmental conditions, revealing an enrichment for missense variants and loci with multiple causal variants. MAGESTIC 3.0 will facilitate the functional analysis of the genome at single-nucleotide resolution and provides a roadmap for improving template-based genome editing systems in other organisms.

## Introduction

Most biological traits are controlled by a complex interplay between an organism’s genotype and its environment. A longstanding promise of biology is that with a deep enough understanding of the molecular mechanisms governing quantitative traits, it should be possible to predict phenotypes from genetic and environmental data. To make progress towards this goal, it is necessary to dissect how each locus in the genome contributes to phenotypic diversity across individuals and species. While association and linkage-based studies have identified thousands of loci impacting quantitative traits, they generally lack the resolution to identify the causal variants in each locus as well as the power to detect rare variants. Hence, the causal variants and molecular mechanisms governing most phenotypic variation in natural populations remain obscure.

Systematic functional screens of individual genetic variants have the potential to overcome the limitations of traditional mapping approaches^1^. Towards this end, we previously developed a high-throughput CRISPR genome editing system based on paired guide RNA/donor DNA templates capable of introducing thousands of genetic variants in parallel in budding yeast termed ***Multiplexed Accurate Genome Editing with Short, Trackable, Integrated Cellular barcodes*** (MAGESTIC)^2^. A key feature of MAGESTIC is a donor recruitment system where a DNA-damage recognizing protein, Fkh1p, fused to the LexA DNA binding domain localizes plasmid donor templates to double-strand breaks to substantially activate homology-directed repair (HDR). Even though donor recruitment substantially increased editing efficiency at individual targets, the overall editing efficiency observed in clones derived from a library of natural variant edits was prohibitively low (∼60%) for effective phenotyping^2^.

To improve the performance of guide-donor plasmid-based systems for variant screens in the present study, we tested MAGESTIC head-to-head against other library-scale guide-donor systems previously developed in yeast, including genetic inactivation of non-homologous end-joining (NHEJ)^3^, single-stranded donor DNA synthesis by bacterial retrons (CRISPEY)^4^ and in vivo assembly of linearized donor plasmids^5^. We assessed editing efficiency, fidelity, and survival (i.e. variant representation), as each can have a major impact on the ability to correctly call phenotypes in large complex libraries. We tested a broad panel of target sites consisting of natural variants across the yeast species both as individual edits to measure efficiency and fidelity and in the context of a pooled library to measure editing toxicity and survival. While donor recruitment provided superior editing overall compared to other approaches, each system showed distinct advantages that could be combined into a single, supercharged donor repair system (MAGESTIC 3.0). MAGESTIC 3.0 proved substantially superior to all previous systems and enabled editing all possible single-nucleotide variants across genomic regions (saturation genome editing). As proof of principle, we used this optimized system to map causal variants in 112 quantitative trait loci and found extensive impact of missense variants on phenotypes, as well as an abundance of loci harboring multiple causal variants.

## Results

### Enhancing homology-directed repair (HDR) for donor-templated CRISPR screens

CRISPR guide-donor libraries enable the parallel introduction thousands of programmed edits into a population of cells. Guide-donor DNA pairs (guide-donors) are first synthesized on oligonucleotide arrays, amplified and cloned into plasmid libraries with unique barcode tags, and finally transformed into an isogenic cell population under conditions such that nearly all transformed cells receive a single plasmid (**Fig 1a**)^2^. The designed edit is introduced by CRISPR-mediated cleavage of the target locus followed by HDR with the donor DNA. The barcode tag on the plasmid specifies the edit and allows for reading out variant function in pooled phenotypic screens by sequencing-based barcode counting of strain abundance (e.g. after competitive growth in diverse environmental conditions).

**Figure 1.**
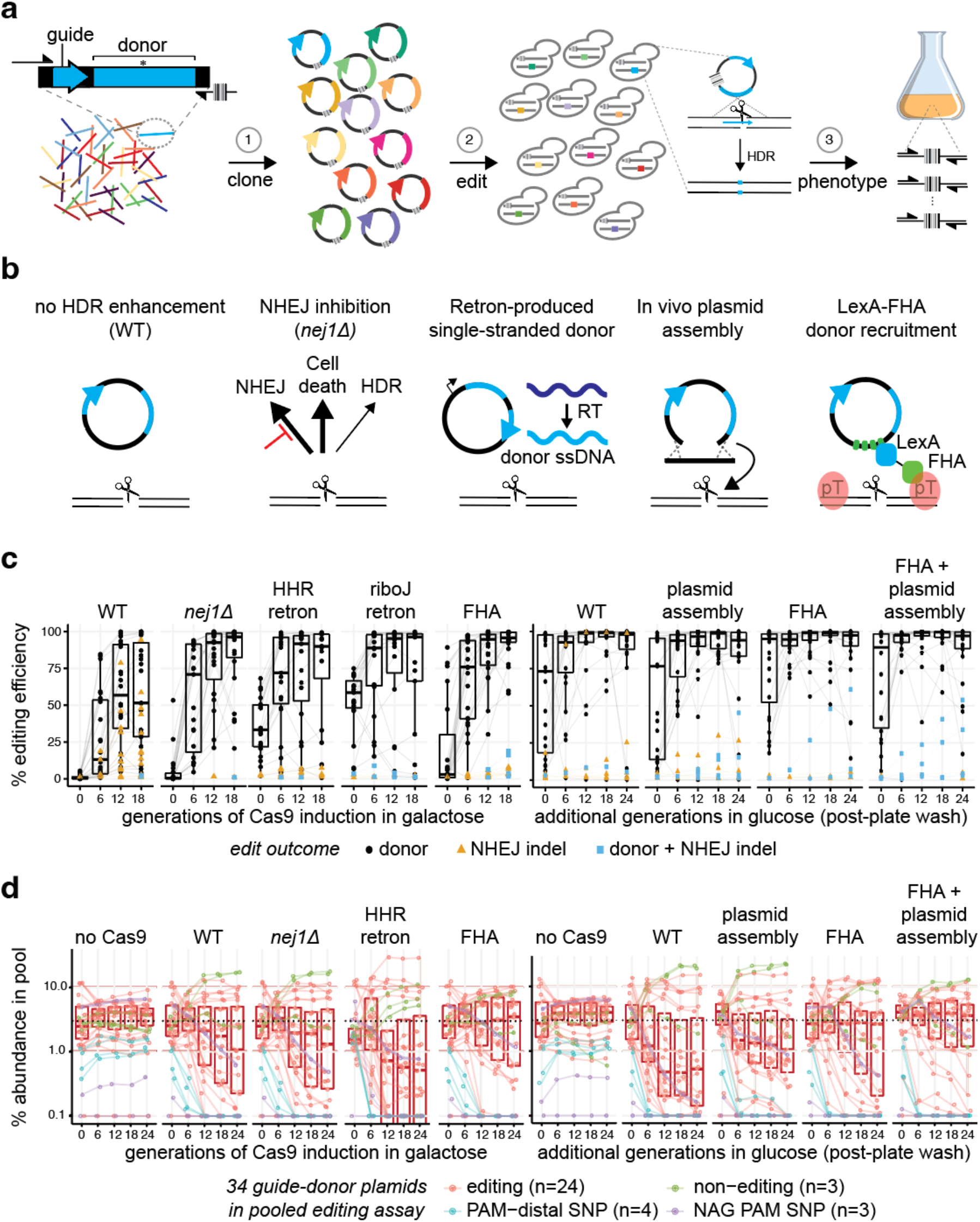
Enhancing homology-directed repair (HDR) with donor DNA templates for CRISPR screens. **a**, CRISPR screens with guide-donor libraries involve (1) oligonucleotide synthesis and barcoded cloning of paired guide RNA/donor DNA repair templates, (2) cell transformation and CRISPR editing by homology-directed repair (HDR), and (3) characterization of growth phenotypes induced by each variant via barcode sequencing (Bar-seq)-based counting of edited strains. **b**, Previous strategies for enhancing editing efficiency used distinct approaches to improve HDR with donor templates. **c**, The editing efficiency for each system is plotted for a panel of 24 natural variant-targeting guide-donors as a function of the number of generations of editing. The HHR retron corresponds to the published CRISPEY system^4^ and the RiboJ retron is a variant of the CRISPEY system developed in Fig. S2. For systems utilizing induction of Cas9 (left half), cells are transferred from non-inducing (glucose) to inducing (galactose) at 0 generations. For editing systems with constitutively expressed Cas9, the 0-generation time point corresponds to editing observed after colony formation on agar plates. For additional generations of editing outgrowth in glucose media, cells were transferred to liquid media after the transformation (right half). For each of the 24 targets, a ∼200 bp region encompassing the edit site was analyzed by on-target NGS. Lines connect the same edit across timepoints. Boxplots show distribution of editing efficiency by donor HDR for all 24 guide-donors (i.e. excluding editing by NHEJ). **d**, Variant abundance during pooled editing. Editing survival was assessed by a competitive growth experiment using a mini-pool consisting of the 24 editing guide-donors (red) assayed in panel c, as well as 3 non-editing cassettes (green), 4 cassettes with PAM-distal SNVs (blue) and 3 cassettes where SNVs result in NAG PAMs (purple). The latter two categories have guides which are expected to cleave the donors at high levels due to tolerance of Cas9 for SNVs at PAM-distal positions and for NAG PAMs. The pool was transformed into (left half) galactose inducible Cas9 systems, induced at time zero by shifting from glucose into galactose, or (right half) constitutive Cas9 systems, where 0g represents a wash of the transformation plate where editing has already begun. The mini pool is constructed initially with all plasmids at near equimolar ratio. Variant ratios were determined at each generation by sequencing of the barcodes in the yeast pools as shown in panel a. Dotted lines indicate 3% abundance. Boxplots include only the 24 editing guides.

A consensus from previous studies developing guide-donor library approaches was that natural HDR efficiencies with plasmid donor DNA are too low for effective screens^2–7^. To enhance efficiency, each study employed a different approach, including genetic inactivation of NHEJ (*nej1Δ*)^3^, in vivo assembly of linearized guide-donor plasmids^5^, recruitment of donor DNA by LexA-Fkh1p (MAGESTIC)^2^, and in vivo production of single-stranded donor DNA with the bacterial Eco1/Ec86 retron system (CRISPEY)^4^ (**Fig 1b**). Each study reported high editing efficiencies (>80-100%) on individual target sites but tested different types of edits at different target sites and measured efficiency via different assays. Therefore, it remains unclear how each of these HDR improvement methods compare with each other on the same set of target sites.

First, we explored whether the MAGESTIC donor recruitment system could be improved. The Fkh1p forkhead-associated (FHA) domain binds phosphorylated threonine residues as part of the DNA damage response and is required for localization to DNA breaks^8^. To test whether it is also sufficient, we constructed a minimal fusion protein containing the LexA DNA binding domain and the FHA domain. This minimal fusion localized to the site of HO-induced breaks by microscopy and gave a substantial, consistent boost in HDR editing over the full-length protein in an editing survival assay at two distinct targets in both haploid and diploid yeast cells (**Supp Fig 1**). We also note that the original MAGESTIC system utilized a tRNA-HDV ribozyme promoter for guide expression, and we showed this promoter led to lower editing efficiency for U-rich guide RNA sequences^2^. We hypothesized these were triggering early termination and lower guide levels and lowering efficiency overall, and therefore switched to the *SNR52* (RNA polymerase III) promoter for this study as it has been shown to be less prone to terminate at stretches of T residues^9^.

Second, we tested whether the retron editing system could be improved through systematic testing of ribozymes flanking the donor-guide cassette, utilizing the same 18mer *ADE2* guide and donor characterized in the CRISPEY study^4^ for consistency (**Supp Fig 2**). We found that the HDV ribozyme was absolutely required on the 3′ end of the guide for detectable editing. Surprisingly, we found that the hammerhead ribozyme (HHR) in the 5′ position employed in the CRISPEY retron system^4^ exhibited poor editing efficiency relative to the 5′ HDV ribozyme **(Supp Fig 2c)**. As a tandem repeat of HDV in both the 5′ and 3′ positions would lead to plasmid instability, we explored whether additional ribozymes or RNA processing elements at the 5′ position could boost editing similarly to HDV. We found that the riboJ ribozyme improved editing efficiency kinetics substantially over HDV, reaching 90% efficiency after 18 generations, compared to 70% for the 5′ HDV and 18% for the 5′ HHR **(Supp Fig 2c)**.

Next, we sought to benchmark each system across a broad panel of 24 natural variants, including guide-donors randomly selected from an *RM11* strain variant library as well as guide-donors which gave rise to unedited clones in the previous MAGESTIC system^2^ (**Supplementary table S1**). With galactose induction of Cas9 and no HDR enhancement (WT), we found a mean HDR editing efficiency of 62%, with unwanted NHEJ indels on 10 targets ranging from 10 to 95% (**Fig 1c**). Both NHEJ inhibition (*nej1Δ*) and the retron systems prevented indel formation and increased HDR editing efficiency, and the riboJ retron overall showed improved editing kinetics compared to the HHR retron, consistent with the earlier results **(Supp Fig 2c)**. Overall, LexA-FHA showed superior editing efficiencies with minimal indel formation in galactose. Next, we analyzed editing with constitutive Cas9 expression in glucose media. We found that editing efficiency from colony transformants on agar plates could be substantially improved with additional liquid outgrowth (**Fig 1c**). Even in the absence of an HDR-enhancing system (WT), 19 guide-donors reached ∼100% donor editing after 18 generations of additional editing in glucose with only 2 targets showing substantial NHEJ indels, suggesting that constitutive editing in glucose is superior to galactose. The indel formation was mitigated by both the linearized plasmid assembly and LexA-FHA. The LexA-FHA donor recruitment system yielded the highest levels of editing efficiency, especially at the earlier stages of editing (**Fig 1c)**.

While editing systems are typically characterized by efficiency and fidelity at individual targets, editing survival is a major factor limiting large-scale CRISPR guide-donor screens^2,5^. In natural variant libraries, there is further the potential for the guide to re-cleave the edited site and/or the donor plasmid since the typical edit (SNV) results in a single mismatch in the guide region which does not always prevent Cas9 cleavage. These issues are important for pooled screens because the ability to measure fitness effects is dependent on the variant starting abundances after editing has gone to completion. To simulate a natural variant library, we pooled together the 24 editing guide-donors with three non-functional guide-donors containing truncated guide scaffolds as controls for no editing toxicity and four with SNVs distal to the protospacer adjacent motif (PAM) at positions 15, 16, 17 and 19 not expected to substantially block cleavage^2^ as controls for high editing toxicity. Additionally, we included three guide-donors generating NAG PAMs, which are expected to be tolerated for Cas9 (re-)cleavage to a variable extent^10^. The 34 guide-donor plasmids were pooled equally and transformed into yeast expressing the different editing systems.

With WT DNA repair, there was a substantial divergence of strain abundance during the editing time course observed with either galactose induction of Cas9 or with constitutively expressed Cas9 in glucose (**Fig 1d**). While NHEJ inhibition and the HHR retron improved editing efficiency, they did not significantly improve variant abundances. Linearized plasmid assembly improved abundances modestly over WT, while LexA-FHA donor recruitment exhibited a strong improvement in overall survival, reducing library skew and enrichment of the non-functional guides considerably compared to the other methods. Strikingly, the combination of donor recruitment and linearized plasmid assembly exhibited an additive effect with improved editing survival compared to either method alone (**Fig 1d**).

Inspection of the editing survival curves for the donor recruitment system revealed three distinct classes of abundance trajectories: those with stable abundances, those with moderate dropout rates, and those with high dropout rates matching the profile of the PAM-distal SNVs (**Supp Fig 4a**). The stable class of guide-donors exhibited similar stability in all systems from 6 generations onwards. Interestingly, however, LexA-FHA appeared to have the greatest benefit at the initial stages of editing outgrowth, where significant skew accumulates the WT and plasmid assembly systems (**Supp Fig 4a**, top row). This is consistent with these edits resistant to cleavage by Cas9. By contrast, the moderate dropout class decreased across all systems at each time point (**Supp Fig 4a**, middle row). Intriguingly, the plasmid assembly method appeared to have the greatest benefit for this class of edits, exhibiting a lower rate of dropout than even LexA-FHA. Strikingly, the combination of donor recruitment and plasmid assembly had beneficial effects on both classes, suggesting that these combining orthogonal HDR enhancement strategies is a promising approach for improving library-scale editing (**Supp Fig 4a**).

Re-examining editing outcomes stratified by abundance curves revealed a strong relationship between NHEJ indel formation and editing toxicity (**Supp Fig 4b**). This was especially apparent in galactose editing in the absence of HDR enhancement. This is consistent with a model where NHEJ indel formation is a minor outcome relative to perfect DSB repair (either through perfect NHEJ^11^ or by homologous recombination with sister chromatids) or cell death^2,12^, but ultimately leads to predominate the editing outcomes of survivors in the absence of HDR enhancement. By contrast, NHEJ indels were only observed to accumulate in 3 and 2 cases with plasmid assembly and LexA-FHA, respectively. These occurred at target sites that had initially edited to 100% efficiency with HDR repair (**Supp Fig 4b**). Overall, these results suggest that higher guide efficacy comes at the cost of increased Cas9 tolerance for mismatches and hence lower SNV edit stability. Furthermore, these data underscore the importance of balancing prolonged editing outgrowth with lower efficiency targets while avoiding excessive re-cutting and library dropout of high-efficiency targets, which is achieved with an outgrowth of 6-12 generations after colony formation (**Fig 1, Supp Fig 4**).

### Combining an improved retron system with donor recruitment

Inspired by the results obtained from combining donor recruitment and plasmid assembly, we revisited the retron system and looked for ways to further improve and integrate it with the donor recruitment approach. To gain insights into how the different ribozymes impact retron donor DNA output, we used an NGS-based approach to simultaneously amplify donor DNA from the single-stranded retron donor as well as the (unedited) target locus in the genome in the absence of Cas9 (**Supp Fig 5**). Surprisingly, the HHR-HDV retron (CRISPEY)^4^ yielded the lowest levels of donor DNA, with less than one donor per genome equivalent (**Supp Fig 5c**). Across all combinations, HHR in the 5′ position consistently lowered retron output independent of the 3′ ribozyme, and HDV in the 3′ position consistently lowered retron output independent of the 5′ ribozyme. This contrasts with the editing results, where the HDV in the 3′ position was required for detectable editing (**Supp Fig 2**). On the other end of the spectrum, the HDV ribozyme in the 5′ position had a dramatic positive effect on retron output, reaching 500-1000 donor ssDNA molecules per cell (**Supp Fig 5**). As the HDV cleaves on its 5′ side and the HHR cleaves on its 3′ side, the 5′ HHR-3′ HDV ribozyme arrangement exposes both ends of the retron transcript to cellular exonucleases which would explain the low donor production. This suggests that the retron transcript benefits from extra sequence on the 5′ and 3′ ends to protect against exonucleases. By contrast, extraneous 5′ or 3′ sequence inhibits guide activity^13,14^ and is unnecessary for stability due to the protective effect of Cas9 binding.

Taken together, the results above suggest that separating the guide RNA from the retron donor and expressing each with optimal flanking elements should improve editing. To test this, we expressed the 5′ HDV retron donor separately from a guide expressed from the *SNR52* promoter and compared editing efficiency to the HHR and RiboJ retron donor-guides from **Supp Fig 2**. To sensitive the system to detect differences in HDR repair and also to simulate lower efficacy guides observed in libraries, we further truncated the guide RNA from an 18mer to a 17mer. We found that the 5′ HDV retron outperformed both the riboJ and HHR retrons, suggesting that additional copies of the retron donor are indeed beneficial for HDR and that the Cas9:guide RNA complex does not need to recruit the retron donor to achieve high editing efficiency, as has been previously suggested^4,15,16^.

Despite the ∼100-fold improvement in retron cDNA output from the HDV retron over the RiboJ retron, this yielded only modest improvements in editing efficiency (see 6, 12 and 18 generation time points, **Fig 2c**). We reasoned that hundreds or even thousands of donor template might not be enough to saturate the edit locus with template in each cell and that template concentration at the target site is limiting. Therefore, we sought to improve the retron further by recruiting it directly to the site of breaks via the LexA-FHA system. We first explored several methods of introducing LexA repeat structures into a retron construct. Introducing two LexA inverted repeats downstream of the donor DNA increased retron production (**Supp Fig 7**) which manifested in improved editing efficiency and survival (**Supp Fig 8**). However, this effect was independent of the expression of LexA-FHA (**Supp Fig 8**) suggesting that the LexA structure itself was enhancing editing simply through improved retron RNA stability rather than recruitment.

**Figure 2.**
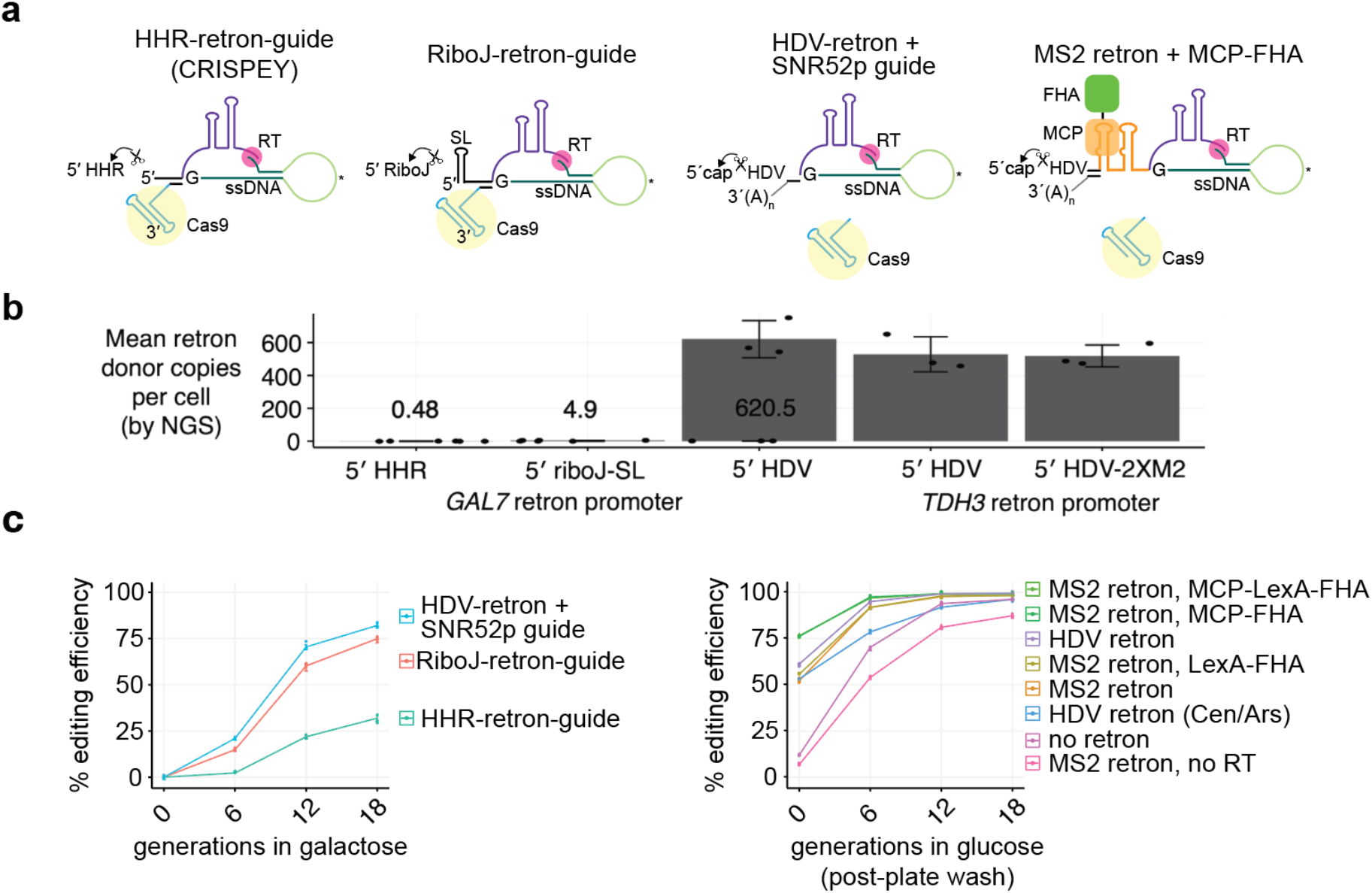
Recruitment of retron donor DNA using the MS2 system and an MCP-FHA fusion protein. **a**, Different arrangement of retrons tested shown from left to right in order of enhanced retron output and editing efficiency. **b**, Retron donor cDNA output from each system as measured in **Supp Fig 5**. The levels for HHR and riboJ retrons are shown above each bar. **C**, Editing efficiency for each retron arrangement shown in **panel a** as a function of generations of Cas9 and retron induction in galactose (left panel) or generations of liquid growth after colony formation on agar plates (right panel). The donor and guide are the same characterized in Sharon et al.^4^ for targeting the yeast *ADE2* gene, where the guide was engineered to have only 18 bp of matching sequence. To give greater sensitivity towards measuring differences in template HDR rates and to simulate the weaker guides which would be observed in a genome-wide library, we artificially weakened this guide with an additional mismatch at position 17. The resulting guide is 5′-cacTTAACGAAATTGCCCCA-3′, where lowercase letters denote mismatches to the target site. All guide-donor plasmids used in the constitutive glucose system are 2μ (high-copy) plasmids, with the exception of the HDV retron (Cen/Ars) shown blue. All systems in the right panel have the guide RNA expressed under the SNR52 promoter, and all have a *TDH3* promoter-driven retron donor except for the “no retron” system, which consists of a donor without any promoter or flanking retron elements. All constitutive systems express Cas9 from the *TEF1* promoter and the RT from the *ADH1* promoter (except for the no RT control), as well as the FHA fusion protein where indicated. Note that the donor transcribed under the retron promoter but without any retron (no RT control) showed reduced editing, suggesting that transcription through the donor is detrimental for plasmid-based template repair.

To explore another means of retron recruitment, we took advantage of the RNA-DNA hybrid nature of the mature retron product as well as fact the 5′ HDV ribozyme remains to the retron transcript after cleavage, protecting the 5′ end of the retron RNA through strong secondary structure. We inserted a tandem repeat of MS2 stem-loop structures in the retron and constructed an MS2 coat protein (MCP)-FHA fusion (**Supp Fig 9**). In the absence of HDR enhancement (no retron), editing at colony formation was only ∼10% (**Fig 2c**). Moving the guide-donor from a single-copy Cen/Ars plasmid to a high-copy 2μ plasmid alone boosted editing efficiency from 53 to 61%. Strikingly, the recruitment of the retron through the MCP-FHA fusion dramatically improved editing efficiency, reaching over 75% at the colony formation stage. Except for the no RT and no retron controls, NHEJ levels in all systems were below 1%, and scaled inversely with improvements in HDR, such that the MCP-FHA fusion showed <0.1% NHEJ indels at the target site (**Supp Fig 8**). While editing with all systems eventually reach close to 100% after 18 generations, the increased donor HDR efficacy results in the need for a shorter editing time course and thus reduced variant skew as shown in **Fig 1** and **Supp Fig 4**. We next tested whether retron recruitment could generate edits by further truncating the *ADE2* guide RNA with mismatches to yield a 16mer, which should lead to little or no cleavage depending on the target site^17^. In this condition, the MS2 retrons yielded 3% editing, while a retron construct with the LexA-LexA stem loop with LexA-FHA expression yielded reduced editing compared to a retron-only control (**Supp Fig 9**). We also tested prime editing^18^ with the same guide lengthened to a full 20mer and the donor edit encoded in the pegRNA tail. This yielded extremely low levels of editing (<1%), suggesting that prime editing is not effective in yeast (**Supp Fig 9**).

### Saturation genome editing with MAGESTIC 3.0

Next, we explored combining all three systems (plasmid donor recruitment by LexA-FHA, ssDNA retron donor recruitment by MCP-FHA, and plasmid assembly) into a single, supercharged editing system termed MAGESTIC 3.0 (**Fig 3a**). To test the ability of MAGESTIC 3.0 to dissect functional natural variants, we challenged it with an assay designed to edit all potential SNVs across genomic regions. We chose ∼200 bp editing windows harboring 6 (SpCas9) or 4 (LbCas12a) non-overlapping guide targets and designed libraries where each guide was paired with a panel of donor DNAs to introduce all 3 SNVs at each position in the 20mer guide sequences and NGG (SpCas9) or TTTV (LbCas12a) PAM sequences (**Fig 3b**). After colony formation, on-target editing outcomes for MAGESTIC 3.0 and each of its sub-systems were assessed by high-throughput sequencing of target region amplicons. In the absence of HDR enhancement, each library exhibited ∼20% editing efficiency. This rose dramatically in the MAGESTIC 3.0 system to 84% for the SpCas9 library and 59% for the LbCas12a library (**Fig 3c**). We note that these libraries harbor differing levels of synthesis errors, with the LbCas12a library exhibiting slightly higher levels with a greater fraction of guides with errors, likely explaining the lower editing efficiency.

**Figure 3.**
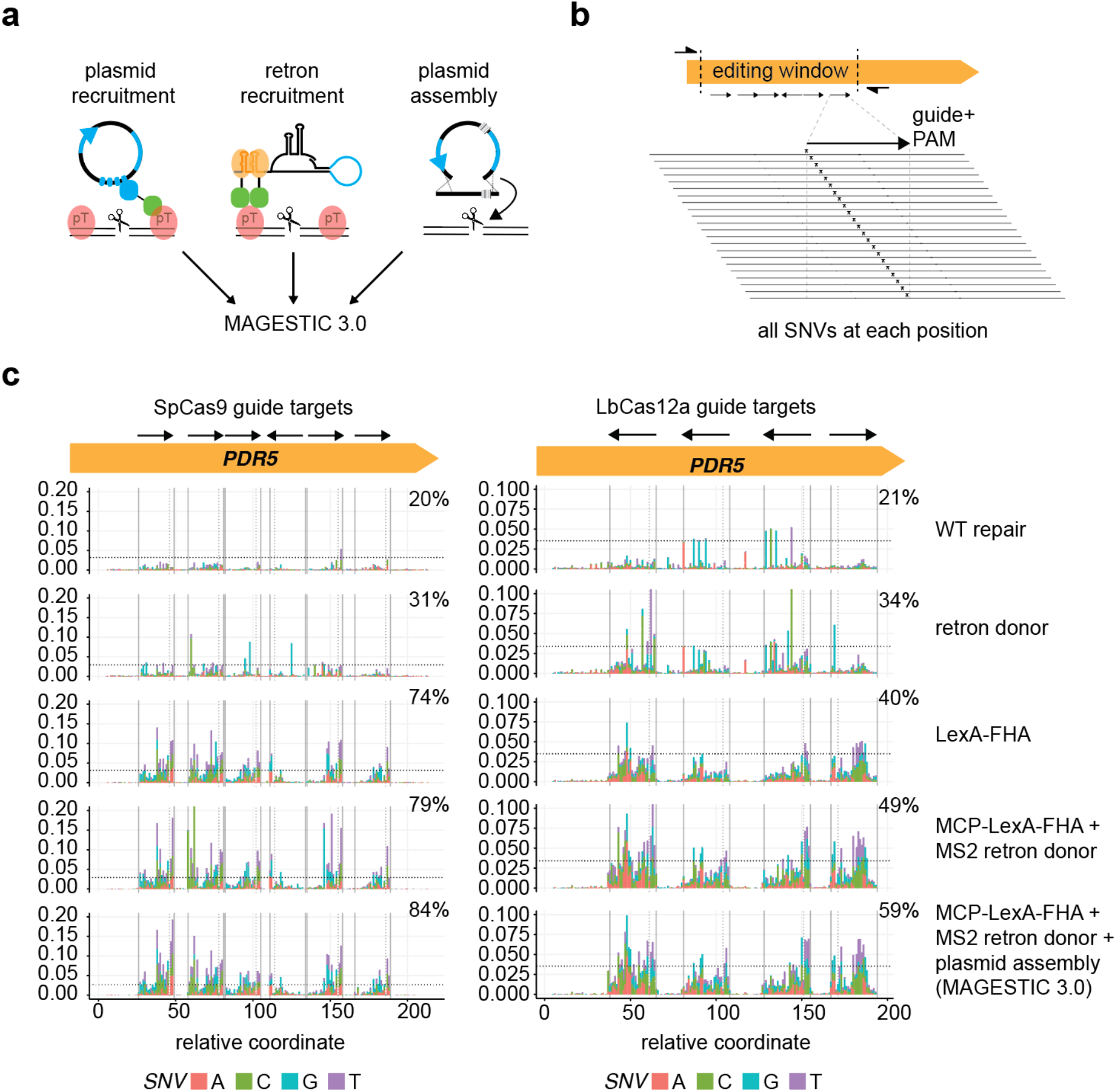
Saturation genome editing with MAGESTIC 3.0. **a**, MAGESTIC 3.0 utilizes three major HDR-enhancing technologies, dsDNA plasmid donor recruitment via LexA sites on the plasmid and the LexA-FHA fusion protein, ssDNA retron donor recruitment via the MS2 system, and plasmid assembly. The use of an MCP-LexA-FHA protein enables simultaneous recruitment of plasmids and retron donor to edit sites. **b**, Editing windows with a panel of non-overlapping guides were selected and all possible SNVs across the guide target region and PAM were engineered into the donor DNAs. **c**, On-target editing rates were quantified by high-throughput sequencing of target region amplicons. The fraction of reads containing SNVs at each coordinate on the x-axis is plotted on the y-axis with stacked bars with the colors representing the SNV introduced. The arrows in the editing window signify the guide and its orientation relative to the target site, with Cas9 guides containing the PAM 3′ of the arrow head (3′ end of the guide), and LbCas12a guides containing the PAM 5′ of the arrow tail (5′ end of the guide).

### Dissecting quantitative trait nucleotides with MAGESTIC 3.0

Finally, we used MAGESTIC 3.0 to edit and phenotype 6,671 variants residing in 112 quantitative trait loci (QTL) previously mapped across the genome for 32 conditions^19^. As each locus contains one or more causal variants hiding amongst dozens to hundreds of potentially benign variants, we used both SpCas9 and a protospacer-adjacent motif (PAM)-relaxed version of LbCas12a to target the greatest possible fraction of variants in each locus. We screened the variant libraries across the same 32 conditions previously tested in the QTL study^19^ by using liquid-based pooled growth assays. We sequenced the barcode composition after 20 generations of competitive growth and calculated log2 fold changes relative to the initial 0-generation time point for each barcode.

Our results recapitulated previously validated causal variants in these loci (*MKT1, PMR1*) and revealed a complex genotype-phenotype map of hundreds of causal natural variants across all loci, with missense variants enriched for effects over non-coding and synonymous variants, as expected (**Fig 4**).

**Figure 4.**
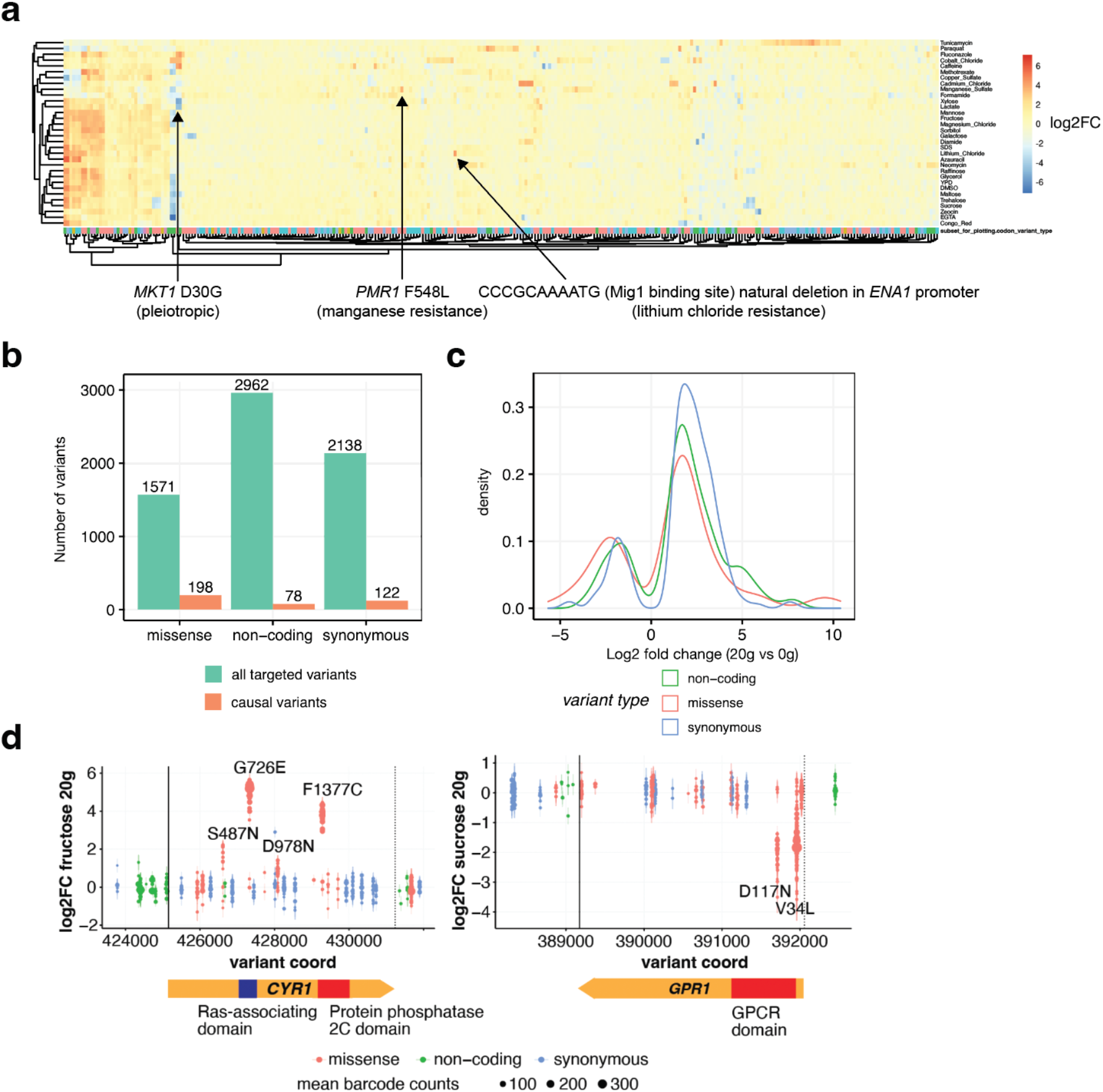
Dissecting causal variants residing in 112 QTL across 32 conditions with MAGESTIC 3.0. **a**, A heat map depicting the log2 fold abundance change for 398 variants which were found to be causal in at least one condition (x-axis), across all 32 conditions (y-axis). Previously reported causal variants in the *MKT1* and *PMR1* genes are highlighted along with a newly uncovered causal variant in the *ENA1* promoter. **b**, Overview of the total number of targeted variants versus the causal variants uncovered in this study stratified by variant type. **c**, The effect sizes for each type of variant are shown as a density plot. Overall missense variants tend to have larger effect sizes than synonymous and non-coding variants. **d**, Log2 fold changes in barcode abundance after 20 generations of growth in fructose (left panel) and sucrose (right panel) for variants in the *CYR1* and *GPR1* genes, respectively. Each point represents an independently edited and barcoded lineage, and type of variant is color-coded. The size of each point corresponds to mean barcode abundance at both 0- and 20-generation time points.

## Discussion

Association-based studies have been highly successful in quantifying the polygenic nature of most traits^20^, yet the ability of these approaches to fully dissect the mechanisms driving traits have major limitations^21^. Two prominent limitations involve linkage disequilibrium, which leads to little or no recombination between individual variants in close proximity on the same haplotype, and rare variants, which do not rise to a high enough frequency to give sufficient statistical power for detecting effects. Systematic perturbation approaches such as CRIPSR screens have potential to address these limitations and provide a major advance forward in our understanding of complex traits^21–24^, by enabling finer-grained dissection of previously mapped QTL and GWAS loci and by discovery of functional variants in previously undetected loci, such as causal variants that are rare or otherwise missed by QTL/GWAS approaches^1,25,26^.

Natural variants pose a significant challenge for CRISPR engineering as the majority are single-nucleotide variants (SNVs). These SNVs must reside within the target sequence of a guide RNA or its protospacer adjacent motif (PAM) to sufficiently block CRISPR cleavage^2,4^. Therefore, guides typically cannot be preselected (e.g. based on predicted efficacy) due to limited PAM availability. Furthermore, the mismatch tolerance of CRISPR nucleases (e.g. SpCas9) can result in repeated cleavage of the donor template and the target site after editing and lead to significant toxicity (i.e. low editing survival) as we demonstrated in this study. In addition, comprehensive profiling of individual genetic variants at the whole-genome scale requires both high efficiency and fidelity of editing as well as scalable and sensitive approaches to characterize phenotypic effects. While some approaches excel in some areas (e.g. high fidelity of SNV editing with prime editing^18^, high efficiency with base editing^27^, they tend to have drawbacks in other areas (e.g. lower editing efficiency with prime editing, restricted edit types and target range with base editing). In this study, we showed that enhanced HDR-based approaches have the potential to solve these problems by simultaneously achieving higher efficiency, fidelity, and superior variant representation. While budding yeast is well known for its predisposition for higher HDR efficiency than most systems, we show that HDR with the donor DNA template is still a major limiting factor in editing performance in the yeast system. Therefore, the development of the MAGESTIC 3.0 system we outline here provides a roadmap for more efficient harnessing of HDR for variant engineering and functional screens in other systems and suggests that adapting the improvements to retron donor DNA production and recruitment of retron donors to DNA breaks outlined in this study will be fruitful in other species and cell lines.

## Material and methods

### Yeast strains

For the comparison of different editing systems, we used the yKR61 strain, a derivative of the DHY214/BY-based wild-type strain where several detrimental alleles have been repaired^28^. For the NHEJ inhibition approach, we knocked out the *NEJ1* gene in yKR61 by utilizing a guide RNA and donor DNA to delete the *NEJ1* open-reading frame to yield yKR139, as previously described^2^. To avoid potential interaction between the *HIS3* marker used in our plasmid assembly approach and the partial HindIII-mediated deletion at the *HIS3* locus present in the BY strain background, we converted the HindIII allele in yKR61 to an entire deletion of the *HIS3* ORF by introduction of a kanMX resistance marker to yield yKR650. For the retron system, we used the ZRS111 strain previously described^4^.

### Natural variant guide-donor plasmids

The guide-donor plasmids assayed in Fig 1 are shown in Supplementary Table S1. The guide-donor cassettes were either amplified from previously isolated MAGESTIC-edited clones or from randomly selected from a *RM11* strain natural variant library^2^ and cloned by Gibson assembly into pKR514, a 2μ (high-copy) plasmid harboring the *SNR52* promoter for guide RNA expression, a 4X tandem array of LexA sites for donor recruitment, and the *FCY1* gene, which is utilized for guide-donor plasmid counterselection after editing. The donors from these plasmids were then used as a template for PCR to generate donor-guide retron constructs for Gibson assembly into either the HHR retron backbone plasmid pKR901, or the RiboJ retron backbone plasmid pKR998. All plasmids were confirmed by Sanger sequencing. The pKR514-based guide donor plasmids were used for all non-retron based editing systems, including the no HDR enhancement control condition. For the plasmid assembly method, the pKR514-based plasmids were cleaved by HindIII, which cuts the *HIS3* ORF in two places. We then amplified a fragment (pF78) spanning these cleavage sites with ∼200 bp of overlap on each side to promote in vivo plasmid assembly. For on-target editing efficiency, each guide-donor plasmid was transformed individually in separate transformations. For glucose editing, cells were plated onto CSM-Ura-His agar plates and incubated at 30°C until colony formation.

In parallel, transformation aliquots were grown in liquid media to facilitate additional liquid outgrowth passages for all targets and editing systems. For galactose editing, transformation aliquots were first grown in CSM-Ura-His liquid glucose media for two passages to select for transformants and establish time zero samples. For the editing time course, 1.5 uL of culture was transferred to 98.5 uL of fresh media each day to allow for 6 generations of growth in 96-well plates. Genomic DNA was prepared from the cultures and the target sites were amplified for high-throughput Illumina sequencing with 2 × 150 bp reads (Novogene). For the editing survival assays, the plasmids were quantified and pooled together at equimolar ratios prior to transformation. Primers were designed to amplify barcoded donor sequences on the plasmid pools to quantify strain abundance by Illumina sequencing with 2 × 150 bp reads (Novogene).

## Data availability

The raw sequencing data reported in this study are available at the SRA database (https://www.ncbi.nlm.nih.gov/sra/) under BioProject accession number PRJNA1067405.

## Code availability

The scripts and codes used for data analysis in this study are available at GitHub (https://github.com/k-roy/MAGESTIC).

## Acknowledgements

This work was supported by NIH grants R01GM121932 and R01HG012446 and an European Research Council (ERC) Advanced Investigator grant (AdG-742804) to L.M.S. K.R.R. was supported by a National Research Council (NRC) postdoctoral associateship. S.C.V. was supported by an Advanced Postdoc Mobility Fellowship (P300PA_177909) from the Swiss National Science Foundation.

