## Supplementary Figures for "Dissecting quantitative trait nucleotides by saturation genome editing"

This file contains:

**Supplementary figures 1 – 9**

**Supplementary references**

### Supplementary figures

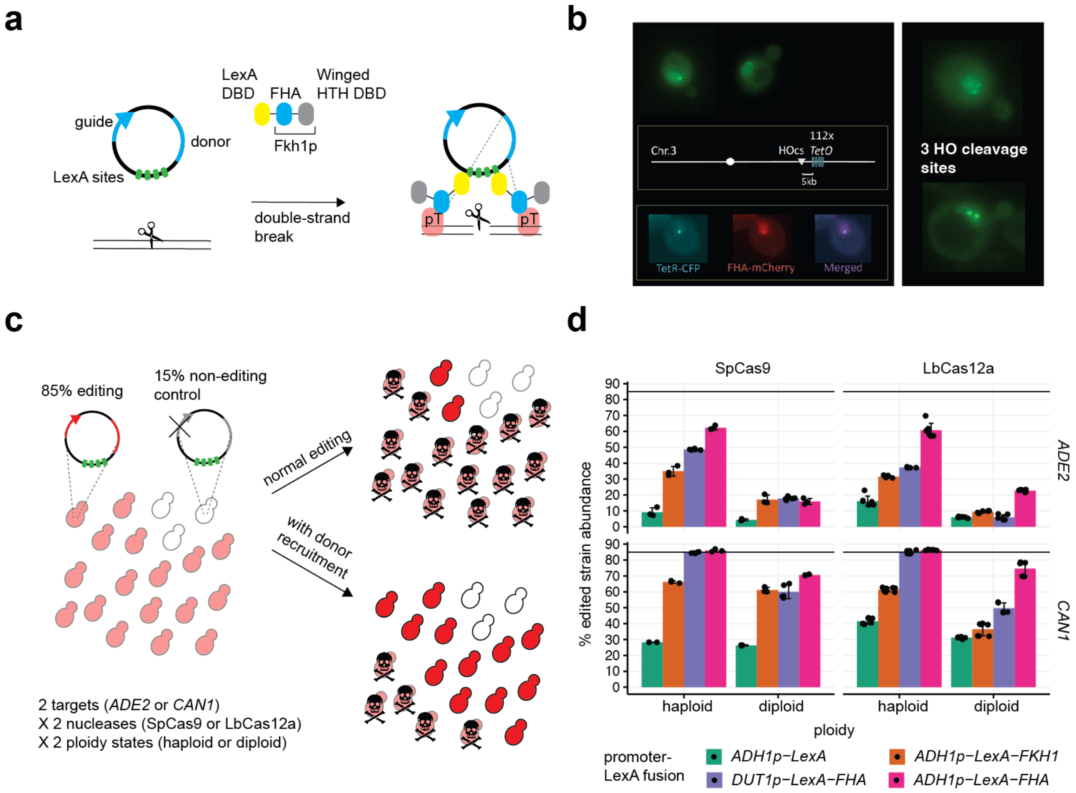


**Supplementary figure 1.** Donor DNA recruitment with the FHA domain of Fkh1p (related to **Fig 1**).

**a,** The donor recruitment protein in the original MAGESTIC system consists of the LexA DNA binding domain (DBD) fused to the yeast Fkh1p protein, which contains a forkhead-associated (FHA) domain and a Winged helix-turn-helix (HTH) DBD.

**b,** Localization of FHA-GFP to a single nuclear focus after induction of a double-strand break by the Homothallic switching (HO) endonuclease (top left). A tandem array of 112 TetO sites was inserted 5 kb downstream of the HO cleavage site at the *MAT* locus on yeast chromosome III to evaluate colocalization of FHA-mCherry and TetR-CFP (left panel). Two additional HO cleavage sites were added at different chromosomal locations to visualize FHA-GFP foci (right panel).

**c,** A spike-in assay for editing survival with guide-donor plasmids targeting two different genes (*ADE2* or *CAN1*) with SpCas9 or LbCas12a in both haploid and diploid yeast cells. The guide-donor pairs utilized in this experiment result in ~100% editing efficiency but low survival when assayed as a single target. The addition of 15% non-editing plasmid simulates the conditions in a complex library where a fraction of the guides can contain synthesis errors or be naturally inefficient. Under normal editing conditions, where efficient guides result in high levels of cell death (top right), non-edited cells become over-represented. Improved HDR increases survival and reduces this skew (lower right). **d,** Amplicon sequencing of the edit sites from washed plates of yeast colonies was used to quantify strain abundance in the populations after editing. 85% edit abundance (solid horizontal line) corresponds to the theoretical maximum of 100% editing survival.

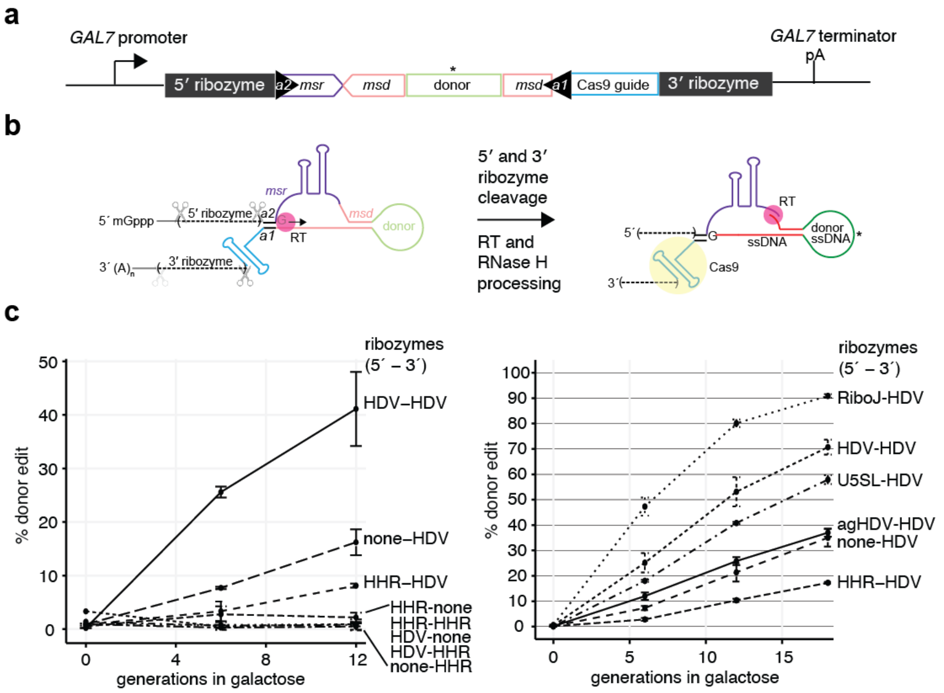


**Supplementary figure 2.** The impact of 5′ and 3′ flanking ribozymes on editing efficiency and retron donor DNA production (related to **Fig 1**).

**a,** Overview of the retron donor-guide system for producing Cas9/gRNA-linked retron donor ssDNA from RNA polymerase II (Pol II) transcripts.

**b,** The location of ribozymes in relation to the 5′ cap (mGppp) and 3′ polyA tail. Depending on whether the ribozymes cleave on their 5′ end or 3′ end, the structured ribozyme can stay bound to the processed retron transcript or be released, exposing the 5′ end of the retron or 3′ end of the guide RNA. After ribozyme cleavage, the retron RNA is reverse transcribed by the RT at a conserved guanosine (G) to create a 2’-5′ RNA-DNA branched structure. The RNA in the RNA-DNA hybrid generated by the RT is degraded by host-cell RNAse H, leaving a looped-out single-stranded donor DNA as a template for HDR.

**c,** Editing efficiency for a panel of ribozymes on the 5′ and 3′ side of the retron donor-guide cassette from Sharon et al.[1] targeting the yeast *ADE2* gene, where the guide was engineered to have only 18 bp of matching sequence. The retron donor introduces CC-to-TG mutation which results in a stop codon. The y-axis indicates the editing efficiency quantified as the % of genomic reads mapping to the designed donor sequence. The x-axis is the number of generations in galactose, which induces Cas9, the RT, and the retron-donor guide transcript. First, we tested all combinations of no ribozyme (none), the hammerhead ribozyme (HHR), and the hepatitis delta virus (HDV) ribozyme (left panel). We then fixed the HDV ribozyme at the 3′ end and tested additional ribozymes and RNA processing elements at the 5′ end, including the anti-genomic HDV (agHDV) ribozyme[2], the U5 snRNA stem-looop cleaved by the yeast RNase III enzyme Rnt1p (U5 Rnt1p SL)[3], and riboJ-SL ribozyme[4], which is a HHR-related ribozyme from the satellite RNA of tobacco ringspot virus (sTRSV) followed by a 23 nucleotide synthetic stem loop (right panel).

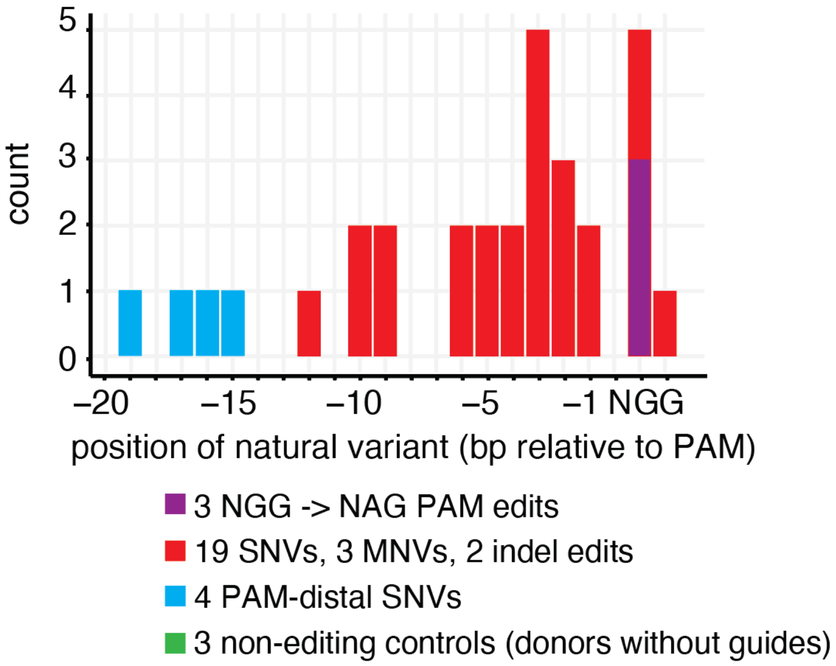


**Supplementary figure 3.** Histogram of PAM-to-edit distances for 34 guide-donors included in the pooled editing survival assay (related to **Fig 1**).

The 24 natural variant editing guides are shown in red. For multi-nucleotide variants (MNVs), the position of the variant closest to the PAM is shown. G to A single-nucleotide variants (SNVs) that generated NAG PAMs are in a separate category (purple) because some of these are expected to be tolerated for additional cleavage by SpCas9, and thus potentially behave like PAM-distal SNPs (cyan). The non-editing control donors (green) are not shown in the histogram as there are no guide RNAs present in their plasmids.

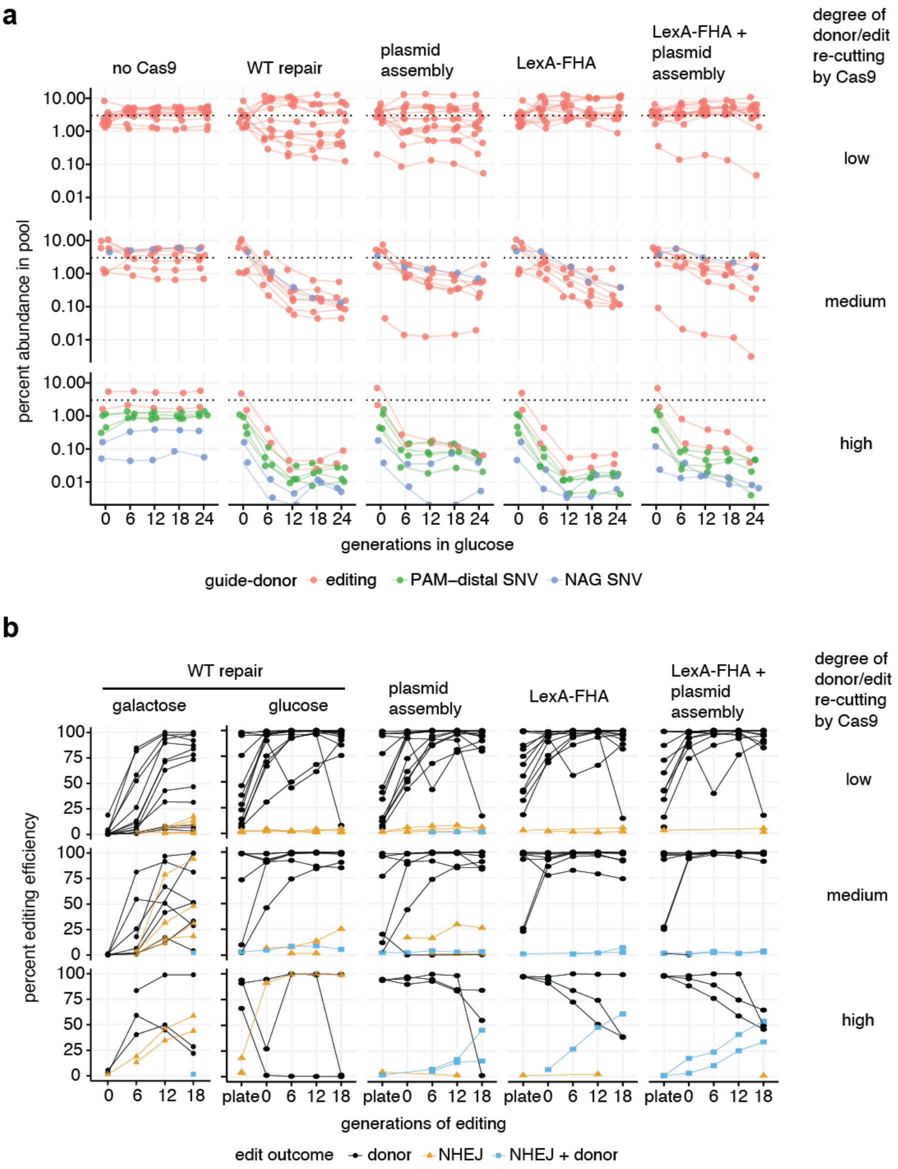


**Supplementary figure 4.** Relationship between editing efficiency and survival based on inferred level of donor cleavage by SpCas9 (related to **Fig 1**).

**a,** Abundance trajectories for the guide-donor plasmids in the pooled competition experiment in Fig 1d in the donor recruitment (LexA-FHA) condition revealed three broad categories of guide-donors: those with stable abundance trajectories, those with moderate decline in abundance, and those with substantial drop out rates similar to the PAM-distal SNVs. Note that one of the NAG PAM SNVs exhibited a moderate decline in abundance and so is plotted in the middle panel. Two of the donors have additional cleavage sites for the HindIII enzyme, which is used to linearize the guide-donor backbones for the plasmid assembly method. Consequently, these start out at substantially reduced levels in the plasmid assembly-based methods. **b,** Abundance trajectories for the guide-donors in different editing systems, with guide-donors stratified by inferred SpCas9 donor/edit cleavage as in panel a. Correct donor templated editing is plotted in black, NHEJ indels are plotted in orange, and donor edits with additional NHEJ indels are plotted in blue.

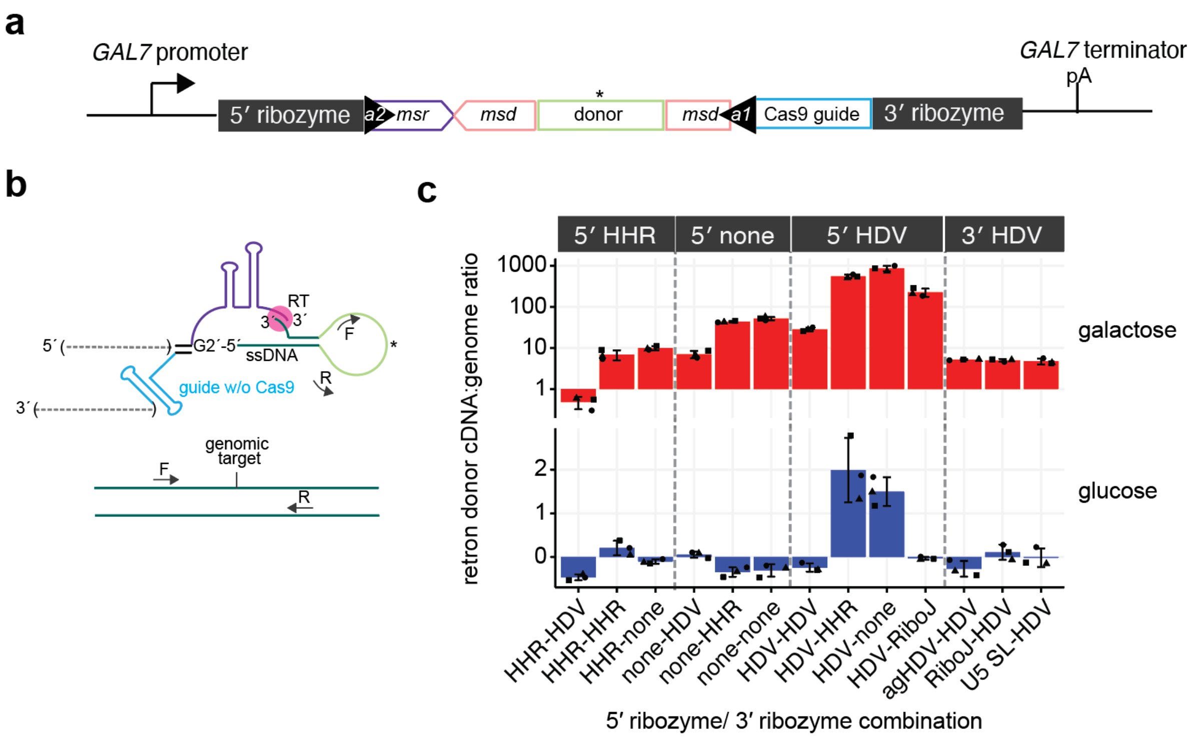


**Supplementary figure 5.** Next-generation sequencing (NGS)-based quantification of retron donor levels (related to **Fig 2**).

**a,** Overview of the retron donor-guide system for producing retron donor ssDNA from RNA polymerase II (Pol II) transcripts. For this experiment, Cas9 was omitted to avoid target site editing.

**b,** Primers are designed to amplify both the single stranded donor template and the genomic target. The donor encodes a CC-to-TG mutation in the middle (asterisk), so the ratio of reads containing the donor mutation relative to the WT genomic locus is proportional to the ratio of donor cDNA PCR template:genome PCR template, with the exception that two strands of the genome are present in each haploid cell (outside of S-phase).

**c,** The ribozyme combinations are grouped according to 5′ ribozymes plotted in **Supp Fig 2c**. Note the y-axis for inducing conditions (galactose) is log10 scale. In glucose, transcription of the retron RT and retron donor are repressed.

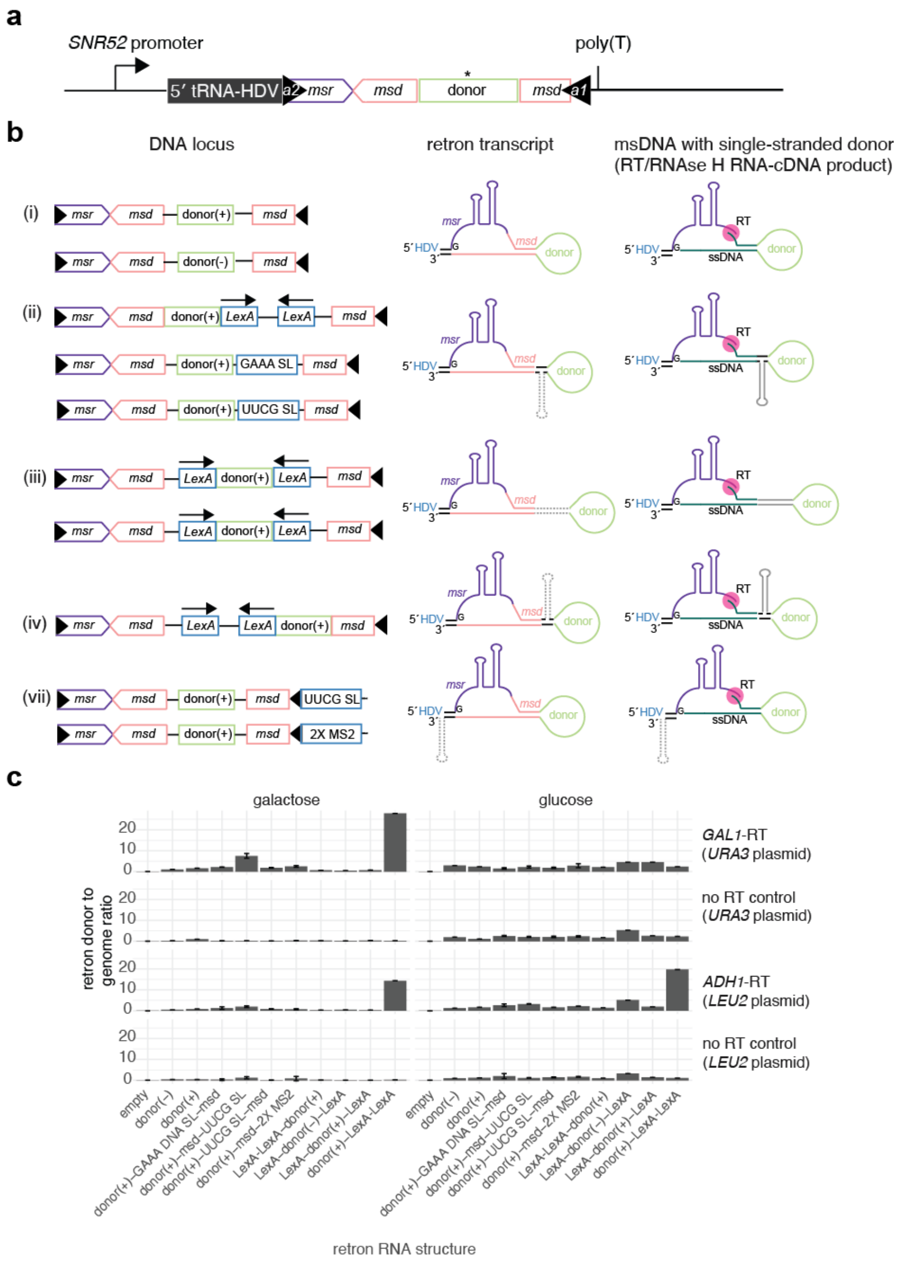


**Supplementary figure 6.** Impact of donor strand and RNA/DNA structures on retron donor production (related to **Fig 2**).

**a,** The retron donor in this experiment was expressed from a fusion promoter consisting of the *SNR52*-tRNA(Tyr)-HDV to stabilize the 5′ end of the retron while keeping it in the nucleus.

**b,** The location of different RNA/DNA structure elements (blue) is shown relative to the terminal inverted repeats (a1 and a2, black triangles), msr (purple), msd (salmon), and donor (green) elements of the retron (left column), with how these structures fold up in the retron RNA (middle column) and msDNA (right column).

**b,** The impact of each of these structures on retron cDNA levels was quantified by NGS. cDNA levels were tested with the RT under the control of the *GAL10* promoter and its no RT control (top two panels), or the *ADH1* promoter and its no RT control (bottom two panels), in either galactose (left) or glucose (right).

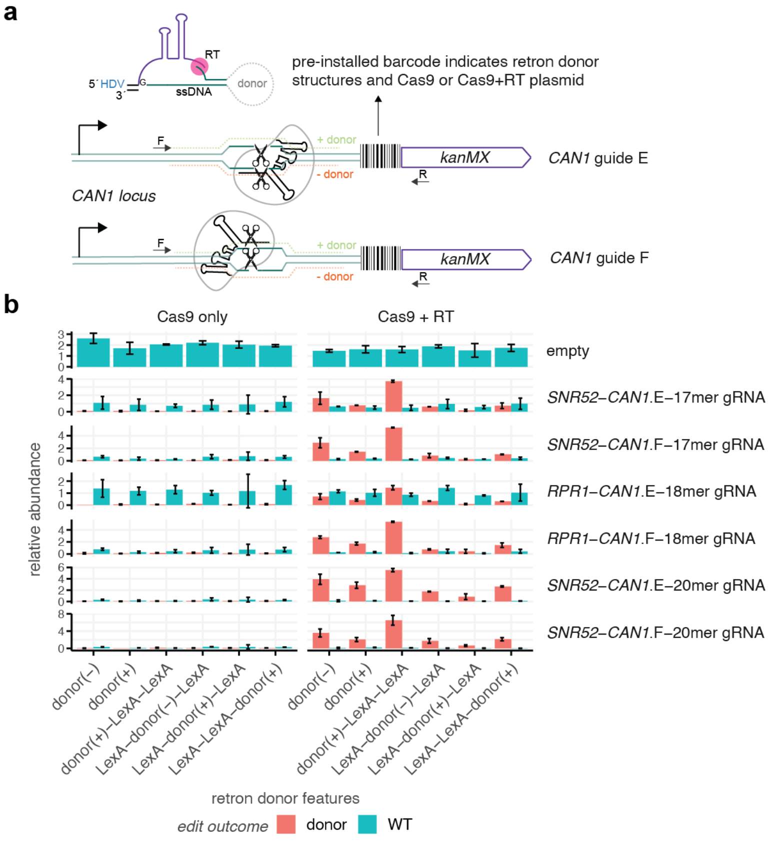


**Supplementary figure 7.** Impact of donor strand and RNA structures on HDR efficacy of retron donor (related to **Fig 2**).

**a**, Various LexA-LexA RNA/DNA structures from Supp Fig 5 were tested in a multiplexed editing experiment where barcodes indicating the retron donor and Cas9 plasmids were inserted adjacent to the target edit site in the *CAN1* gene. To test potential relationships between guide RNA strand and retron donor strand, two overlapping guide RNA target sites (denoted E and F) were selected to have the same predicted SpCas9 cleavage site but with binding to opposite strands (sequenced are 5′-TCACAAACACACCACAGACG-3′ and 5′-ATGGTATTGACCCACGTCTG-3′, respectively). A plasmid harboring each LexA structure variant along with either Cas9 only (left) or Cas9 + RT (right) were transformed into three separate barcoded strains. This approach allowed all strains to be pooled equally prior to transforming a third plasmid containing only the guide RNA. In each NGS read, both the barcode and edit site are sequenced, enabling quantitation of both editing efficiency (ratio of donor to WT on y-axis) and editing survival (total abundance on y-axis). Cas9 and RT were expressed from the constitutive *TEF1* and *ADH1* promoters, respectively. To simulate weaker guides, mismatches were introduced at positions 20, 19, and 18 for the guides expressed from the constitutive *SNR52* promoter. The *RPR1* promoter generates a stable leader sequence[5] on the guide RNA while the *SNR52* leader is efficiently processed by yeast RNase III Rnt1p[6]. The fact that the *RPR1* promoter on 18mer guides shows lower efficiency than the *SNR52* on 17mer guides clearly demonstrates the negative impact of extraneous 5′ sequence on guide activity.

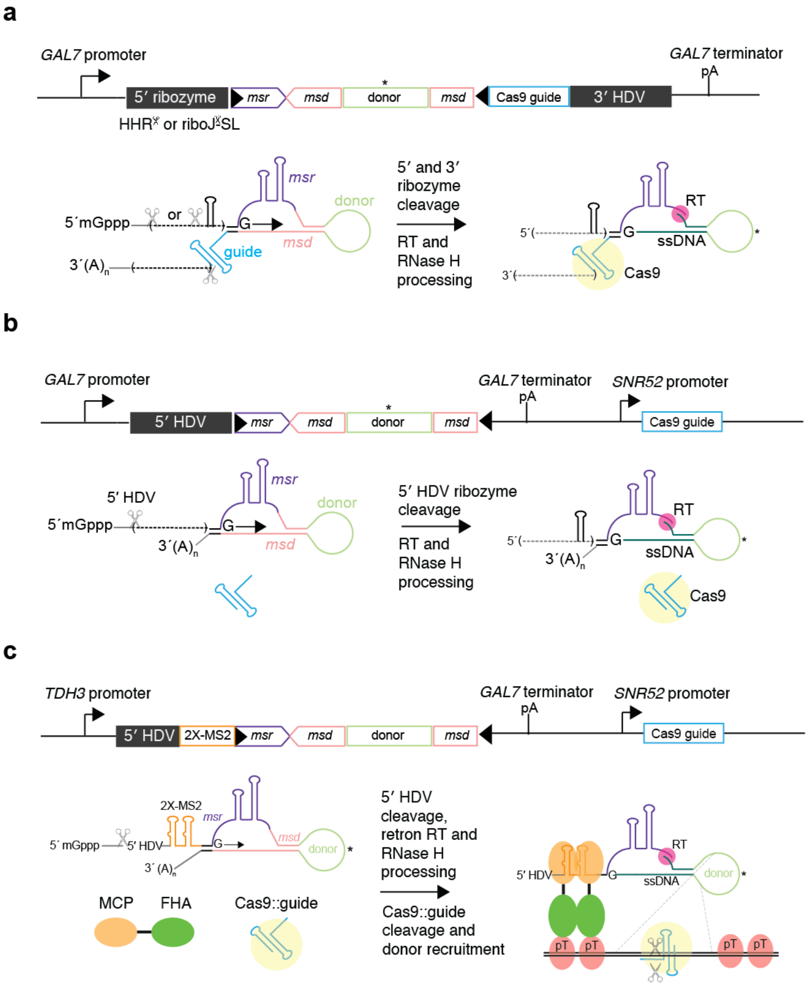


**Supplementary figure 8.** Overview of different retron donor systems, expressed either as a fusion with the guides (a) or with the guides expressed separately from the SNR52 promoter (b, c). Panel c shows the MS2-based donor recruitment system (related to **Fig 2**).

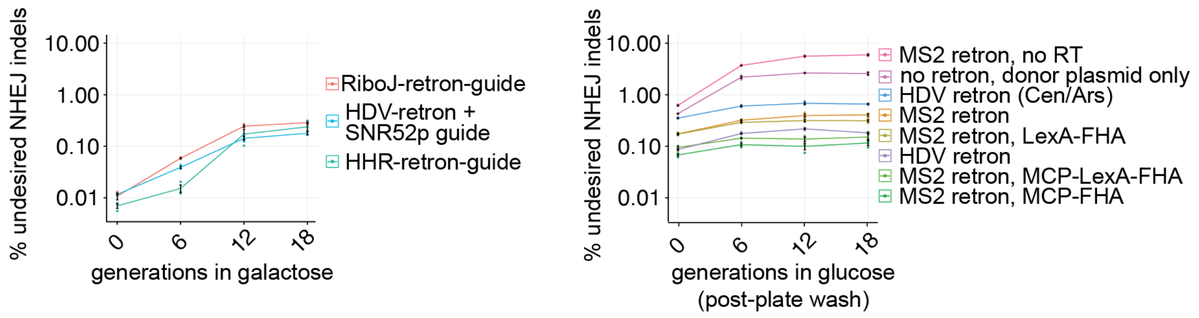


**Supplementary figure 8.** NHEJ indel levels generated by different retron editing systems (related to **Fig 2**).

The same dataset from Fig 2 was analyzed for NHEJ indel levels in the vicinity of the cut site. All functional retron systems result in indel levels below 1%. Note the inverse correlation between donor editing efficiency in Fig 2 and NHEJ indel formation shown here. The systems with the highest donor editing efficiency (HDV-retron + SNR52p guide for galactose; MS2 retron, MCP-FHA and MCP-LexA-FHA for glucose) exhibit the lowest NHEJ levels.

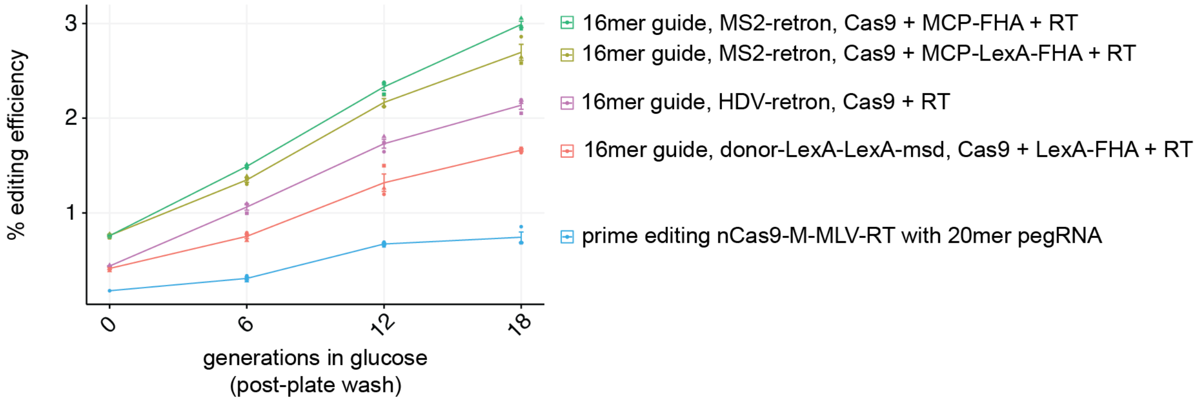


**Supplementary figure 9.** Editing performance with substantially weakened guide RNAs versus prime editing (related to **Fig 2**).

The same guide utilized in Fig 2 was further reduced in efficacy by introducing an additional mismatch at position 17 from the PAM, effectively turning it into a 16mer with the sequence 5′-cacgTAACGAAATTGCCCCA-3′. The best performing systems from **Fig 2** (MS2 retron, MCP-FHA and MCP-LexA-FHA) were tested against the HDV retron donor or a retron donor-LexA-LexA construct, where the LexA stem loops have potential to form a double-stranded stem loop in the retron ssDNA as shown in **Supp Fig 5**. These LexA sites would allow for donor recruitment either directly from the plasmid or via the retron ssDNA. For a comparison with prime editing, the same guide was lengthened to a fully matching 20mer with the pegRNA sequence 5′-TTATTAACGAAATTGCCCCAgtttaagagctatgctggaaacagcatagcaagtttaaataaggctagtccgttatcaacttgaaaaagtggcaccgagtcggtgcTTGTGAGGCCTTcaGGCAATTTCGTTA-3′, where the long stretch of lowercase letters indicates the sgRNA scaffold and the lowercase ‘ca’ represents the edit encoded in the pegRNA 3′ tail.
